# Harnessing Global HLA Data for Enhanced Patient Matching in iPSC Haplobanks

**DOI:** 10.1101/2024.10.15.618361

**Authors:** Martin Maiers, Stephen Sullivan, Christopher McClain, Christina Leonhard-Melief, Marc L. Turner, David Turner

**Author notes:** **Corresponding author:** Martin Maiers, National Marrow Donor Program, 500 N. 5^th^ St., Minneapolis, MN 55401, USA.

## Abstract

Several countries have either developed or are developing national induced pluripotent stem cell (iPSC) banks of cell lines derived from donors with HLA homozygous genotypes (two identical haplotypes) prevalent in their local populations to provide immune matched tissues and cells to support regenerative medicine therapies. This ‘haplobank’ approach relies on knowledge of the HLA genotypes of the population to identify the most beneficial haplotypes for patient coverage, and ultimately identify donors or cord blood units carrying two copies of the target haplotype. A potentially more efficient alternative to a national bank approach is to assess the haplotypes required to provide global patient coverage and to produce a single, global haplobank. Toward that end, we have developed an algorithm to prioritize HLA haplotypes that optimize coverage across the global population.

We analyzed data from eighteen countries participating in the Global Alliance for iPS Therapy (GAiT). A representative pool of HLA genotypes, reflecting the HLA of patients, was derived by sampling from each country’s WMDA hematopoietic stem cell donor registry, or surrogate population. An algorithm was created based on HLA-A, -B and -DRB1 haplotype homozygous types with population HLA matching coverage defined by the absence of Host versus Graft (HvG) mismatches at these loci. HLA matching coverage was determined by iteratively selecting HLA haplotypes that provide the largest coverage against patient HLA genotypes sampled from the same population, excluding genotypes compatible with previous iterations. The top 10 haplotypes for each of the 18 countries were identified with patient coverage ranging from 19.5% in Brazil to 63.8% in Japan, with a mean coverage of 33.3%. In a ‘global’ model, utilizing the 180 most frequent haplotypes across all 18 populations (equivalent to 10 lines per country), the patient coverage ranged from 54.6% in India to 81.7% in Sweden, with a mean of 68.4%. Our findings demonstrate that global collaboration could more than double the potential for patient HLA matching coverage.

Interrogation of unrelated hematopoietic stem cell donor registry and cord blood bank HLA data demonstrated that HLA-A, -B, and -DRB1 homozygous donors for the top 180 global haplotypes are widely available. These results show that a globally coordinated strategy for haplobanking would reduce redundancy and allow more patients to be treated with the same investment.

**Visual Abstract:** 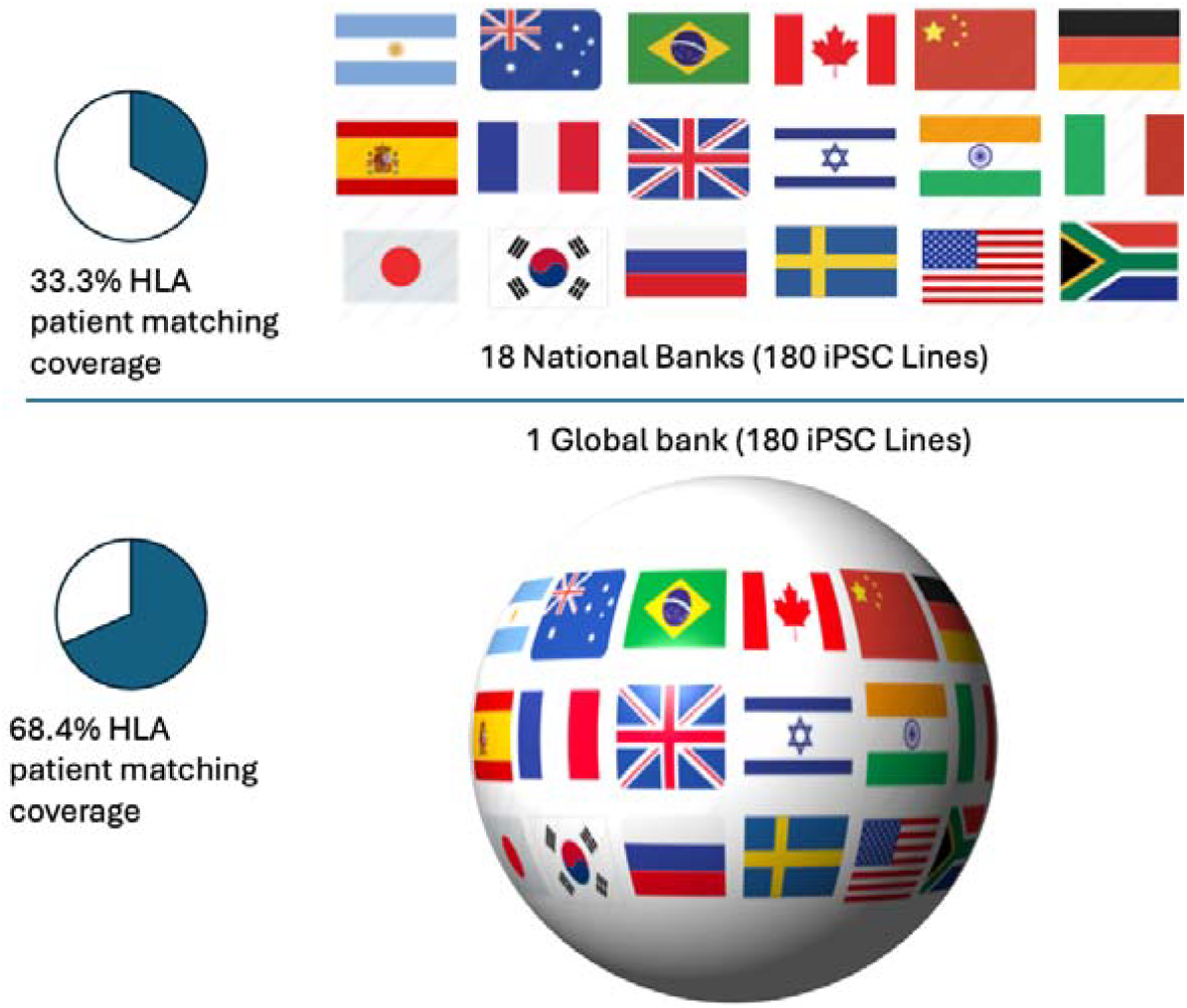

**Highlights:**

- iPSC banks are being developed targeting HLA coverage to meet national needs
- Global collaboration has the potential to substantially increase the efficiency of this effort

## Introduction

Cell and tissue therapies derived from induced pluripotent stem cells (iPSC) are a potential future treatment for many degenerative and chronic conditions, but it is recognized that the recipient immune response represents a major hurdle to the introduction of such therapies (Fairchild et al. 2016; Petrus-Reurer et al. 2021; Solomon, Pitossi, and Rao 2015). Autologous iPSC overcome this concern and most early clinical trials made use of autologous and embryonic stem cell (ESC) derived treatments (Crow 2019). However, for more widespread use these approaches may not be feasible due to cost, time for production, ethical considerations and complexity (Doss and Sachinidis 2019). For allogeneic cells, an assessment of clinical studies initiated since 2018 shows that iPSC now predominates over ESC (Kobold et al. 2023). For further development of the use of allogeneic iPSC, two broad approaches for the mitigation of the patient immune response are advocated; using banked iPSC cell lines representing common HLA types to enable some degree of matching with patients (haplobanking) (Sullivan, Fairchild, et al. 2020), or the creation of hypoimmunogenic lines where the expression of immune stimulating molecules have been down regulated or eliminated (Han et al. 2019).

The former method has been progressed in several countries with the creation of iPSC haplobanks (Álvarez-Palomo et al. 2021; Yoshida et al. 2023; Tian et al. 2022; Lee et al. 2018; Taylor et al. 2005). Cells derived from HLA matched iPSC lines have been used in early safety trails for macular degeneration, and grafts survived for a one-year period in the absence of immunosuppression (Sugita et al. 2020). Although these strategies for overcoming immune activation are not mutually exclusive (Xu et al. 2019), given the necessity for regulatory approval for products derived from iPSC lines, it may be that methods using HLA matched cell lines will pose fewer safety concerns and therefore will face fewer regulatory challenges than those involving genetic manipulation. Other immunological barriers will also need to be considered even if full HLA matching can be achieved using iPSC from haplobanked donors. Some of the concerns around activation of NK cells and minor histocompatibility antigens have been investigated in animal models, with evidence of both NK cell and T cell activation even when full MHC matching is present (Kawamura et al. 2016; Morizane et al. 2017). It is therefore possible that recipients of allogeneic HLA matched products will still need some degree of immunosuppression to limit immunological activation.

Haplobanking requires the identification of donors who are homozygous for HLA haplotypes, thereby allowing matching to patients who have a genotype with at least one shared allele at each locus (Gourraud et al. 2012). HLA polymorphism varies extensively geographically, therefore different haplotypes are required for coverage of different populations (Pappas et al. 2015). Numerous studies have investigated common HLA haplotypes in local populations to identify donors that should be used to create local cell line banks (Nakatsuji, Nakajima, and Tokunaga 2008; Taylor et al. 2012; Lee et al. 2018; Belen Alvarez-Palomo et al. 2022), aiming to cover as high a percentage of the population with HLA matched donor material with the least number of iPSC cell lines. This is easier in some areas because of more limited HLA variation but still requires many cell lines in each population to reach coverage that might be deemed acceptable from an equity of access to healthcare perspective. More recently (Turner et al. 2013; Sullivan, Ginty, et al. 2020; Wilmut et al. 2015) there has been growing recognition that, with access to appropriate HLA haplotype frequency data, modeling could be performed at the global level and with an understanding of the HLA overlap between populations these national efforts could be targeted in a way that strives for a global optimum of patient coverage i.e. harnessing local efforts to produce iPSC with a view to providing the greatest patient HLA matching coverage on an international basis.

The Global Alliance for iPSC Therapies (GAiT) (Sullivan, Ginty, et al. 2020) is a non-profit organization established in 2013 with a mission “to enable the global human community the opportunity to benefit from the new generation of cell therapies by facilitating the development of, and access to, clinical-grade and haplotyped induced pluripotent stem cells for the manufacture of cell therapy products”. With the support of its international partners, GAiT already has an early position on manufacturing, regulatory and quality standards (Sullivan S 2018, Kim JH 2019). Another of GAiT’s main objectives is to model the requirements for an international bank of iPSCs from individuals who are homozygous for HLA haplotypes using high resolution HLA typing data from haematopoietic stem cell registries. It is hoped that this assessment will identify key HLA haplotypes that could reduce the number of donors required globally to provide HLA matched iPSC derived therapies for the greatest number of patients. In this study, we sought to achieve this goal by mobilizing the vast resource of the global network of haematopoietic stem cell registries.

## Material and Methods

### Genotype data

HLA genotypes were sampled from the WMDA registries of the countries included in the analysis (Figure 1 and Table 1). Haplotype frequencies were derived from the data using a maximum likelihood method (Israeli et al. 2021). Inclusion criteria required DNA-based typing at HLA-A, -B and -DRB1. Whenever possible, this was further restricted to donors that were typed at these loci with DNA-based methods at the time of recruitment to avoid bias. These frequencies were used to impute the HLA genotypes (Israeli, Gragert, et al. 2023) and the posterior distribution was randomly sampled to obtain an unambiguous multi-locus genotype for each individual following a method similar to that used in previous studies (Jonna Clancy et al. 2022). For coverage analysis, 100,000 individuals were randomly sampled, or the entire registry population was used for registries with fewer than 100,000 donors.

**Table 1.** Donors and Cord Blood Units considered as potential starting material for this study (based on WMDA inventory as of 2021-07-15)

| # | Country | ISO-3166:2 code | Number of Donors | Number of Cord Blood Units | Total |
| --- | --- | --- | --- | --- | --- |
| 1 | Argentina | AR | 264,741 | 4,093 | 268,834 |
| 2 | Australia | AU | 176,616 | 37,487 | 214,103 |
| 3 | Brazil | BR | 5,019,313 | 17,805 | 5,037,118 |
| 4 | Canada | CA | 494,622 | 16,425 | 511,047 |
| 5 | China | CN | 1,458,544 | 80,380 | 1,538,924 |
| 6 | Germany | DE | 9,786,284 | 38,836 | 9,825,120 |
| 7 | Spain | ES | 459,311 | 62,987 | 522,298 |
| 8 | France | FR | 346,757 | 38,154 | 384,911 |
| 9 | UK | GB | 2,243,849 | 27,879 | 2,271,728 |
| 10 | Israel | IL | 1,373,259 | 16,765 | 1,390,024 |
| 11 | India | IN | 590,418 | 0 | 590,418 |
| 12 | Italy | IT | 474,283 | 39,030 | 513,313 |
| 13 | Japan | JP | 536,259 | 0 | 536,259 |
| 14 | S. Korea | KR | 361,204 | 55,076 | 416,280 |
| 15 | Russia | RU | 127,742 | 6,489 | 134,231 |
| 16 | Sweden | SE | 214,790 | 4,419 | 219,209 |
| 17 | USA | US | 9,436,116 | 272,356 | 9,708,472 |
| 18 | South Africa | ZA | 72,652 | 0 | 72,652 |
|  | Total |  | 33,436,760 | 718,181 | 34,154,941 |

**Figure 1.**
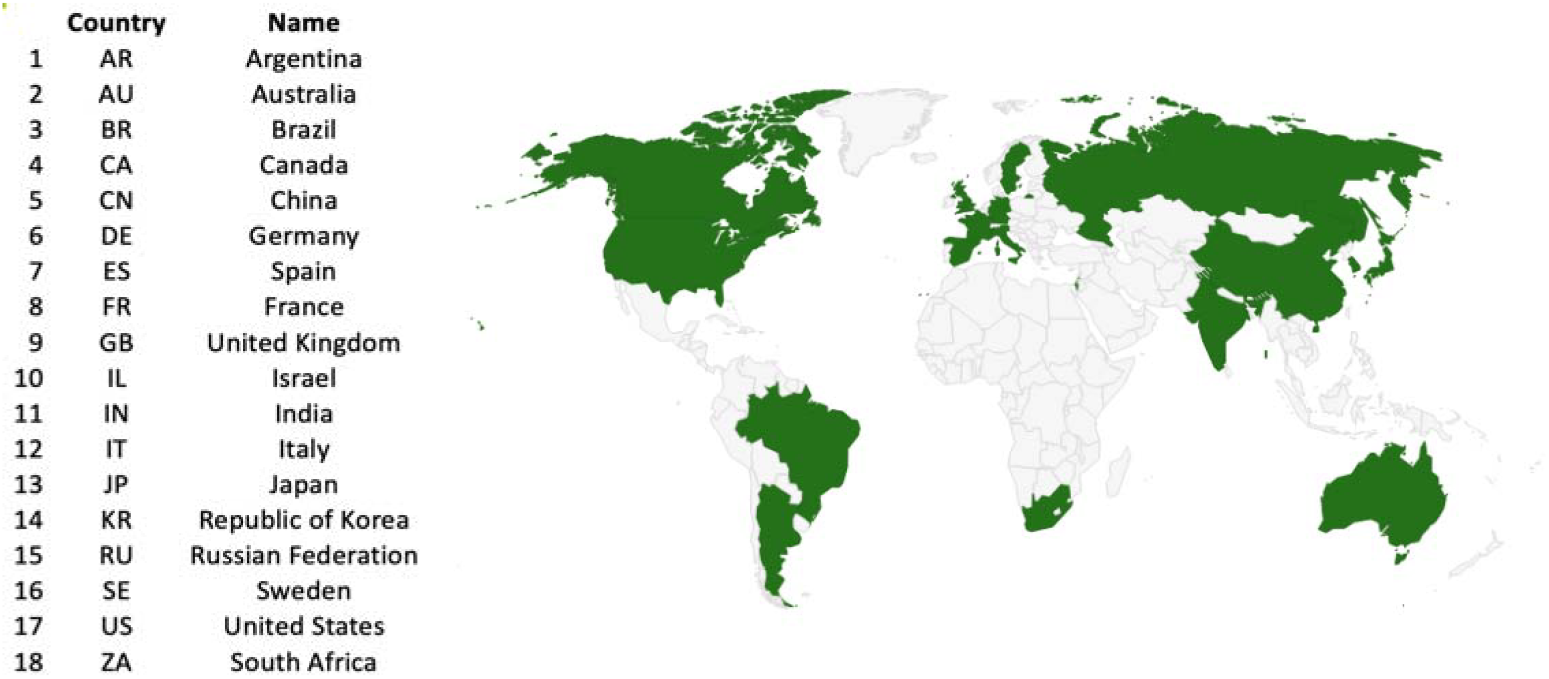
Eighteen countries included in the analysis based on membership in GAiT.

The HLA was reduced to 2-field alleles collapsed over the antigen recognition domain (ARD) defined as exon 2 for class II and exons 2 & 3 for class I. The genotypes were reduced to 3-locus (A, B & DRB1) and the homozygosity status was noted.

### Greedy Coverage algorithm

The national local greedy coverage analysis was performed using a greedy algorithm (Israeli, Krakow, et al. 2023) based on a sample of 100,000 donors with complete typing for HLA-A, -B and -DRB1 (note that for Russia and South Africa smaller samples were used). This was done to avoid bias due to the variation in sample size of the 18 donor registries. An initial pool of candidate homozygous genotypes was generated from the sample, requiring at least one in the donor pool. Each candidate homozygous donor genotype was evaluated for its coverage of the patient pool, defined as the total number of the sample sharing at least one allele at HLA-A, -B and -DRB1 with the target genotype. The genotype with the highest coverage value was selected and all patients covered by this genotype were removed from subsequent iterations. This process was repeated, with the remaining patient pool used to evaluate the remaining candidate homozygous donor genotypes, until the number of cycles was completed or the pool of homozygous donor genotypes was exhausted.

The global model followed a similar approach but included the candidate homozygous donor pool from all 18 countries. Patient coverage was assessed on a per-population basis, with the highest coverage genotype being the one that provided the greatest average coverage across all populations. As with the national models, covered patients were removed after each iteration, and the process continued until either the cycle limit was reached or the pool of homozygous donor genotypes was exhausted.

### Statistical Analysis

The PCA analysis was performed using R (version 4.3.1), *prcomp* and plotted using the *ggfortify* library. The 2D clustering was performed using R, *heatmap*.

## Results

Independent coverage models were developed for 18 countries based on their national HLA genotype distribution as modeled by their donor and CBU pool. Each model was optimized by selecting haplotypes iteratively based on covering the largest proportion of the remaining patient pool. Each national model ran for 10 and 30 iterations. The full tables are in Supplemental Material 1. Using 10 iterations the cumulative coverage ranged from 48.0% for Japan to 19.5% for Brazil (Figure 2), with an average of 33.3% and with 30 iterations from 63.8% for Japan to 32.1% for Brazil, average 47.7% (Table 2).

**Table 2.** Percentage patient coverage for locally optimized top 10 or 30 haplotypes and optimized global 180 haplotypes.

| Country | Coverage by top 10 local haplotypes (%) | Coverage by top 30 local haplotypes (%) | Coverage by top 180 global haplotypes (%) | Percentage point Improvement of global 180 over local 10 | Percentage point Improvement of global 180 over local 30 |
| --- | --- | --- | --- | --- | --- |
| **Sweden** | 47.6 | 62.9 | 81.7 | 34.1 | 18.8 |
| **UK** | 44.1 | 58.0 | 79.0 | 34.9 | 21 |
| **Germany** | 38.3 | 55.5 | 79.0 | 40.7 | 23.5 |
| **Australia** | 39.9 | 53.6 | 77.5 | 37.6 | 23.9 |
| **Russia** | 34.1 | 40.1 | 76.5 | 42.4 | 36.4 |
| **France** | 31.8 | 44.6 | 73.6 | 41.8 | 29 |
| **Canada** | 30.9 | 45.0 | 72.2 | 41.3 | 27.2 |
| **Spain** | 30.7 | 47.0 | 70.3 | 39.6 | 23.3 |
| **Korea** | 40.0 | 57.8 | 68.8 | 28.8 | 11 |
| **Japan** | 48.0 | 63.8 | 67.6 | 19.6 | 3.8 |
| **USA** | 29.1 | 42.3 | 67.1 | 38 | 24.8 |
| **Italy** | 25.6 | 40.8 | 66.7 | 41.1 | 25.9 |
| **China** | 34.2 | 52.3 | 62.3 | 28.1 | 10 |
| **Israel** | 28.5 | 48.3 | 61.6 | 33.1 | 13.3 |
| **South Africa** | 27.0 | 32.8 | 59.6 | 32.6 | 26.8 |
| **Argentina** | 21.8 | 35.3 | 58.6 | 36.8 | 23.3 |
| **Brazil** | 19.5 | 32.1 | 55.3 | 35.8 | 23.2 |
| **India** | 28.9 | 46.3 | 54.6 | 25.7 | 8.3 |

**Figure 2.**
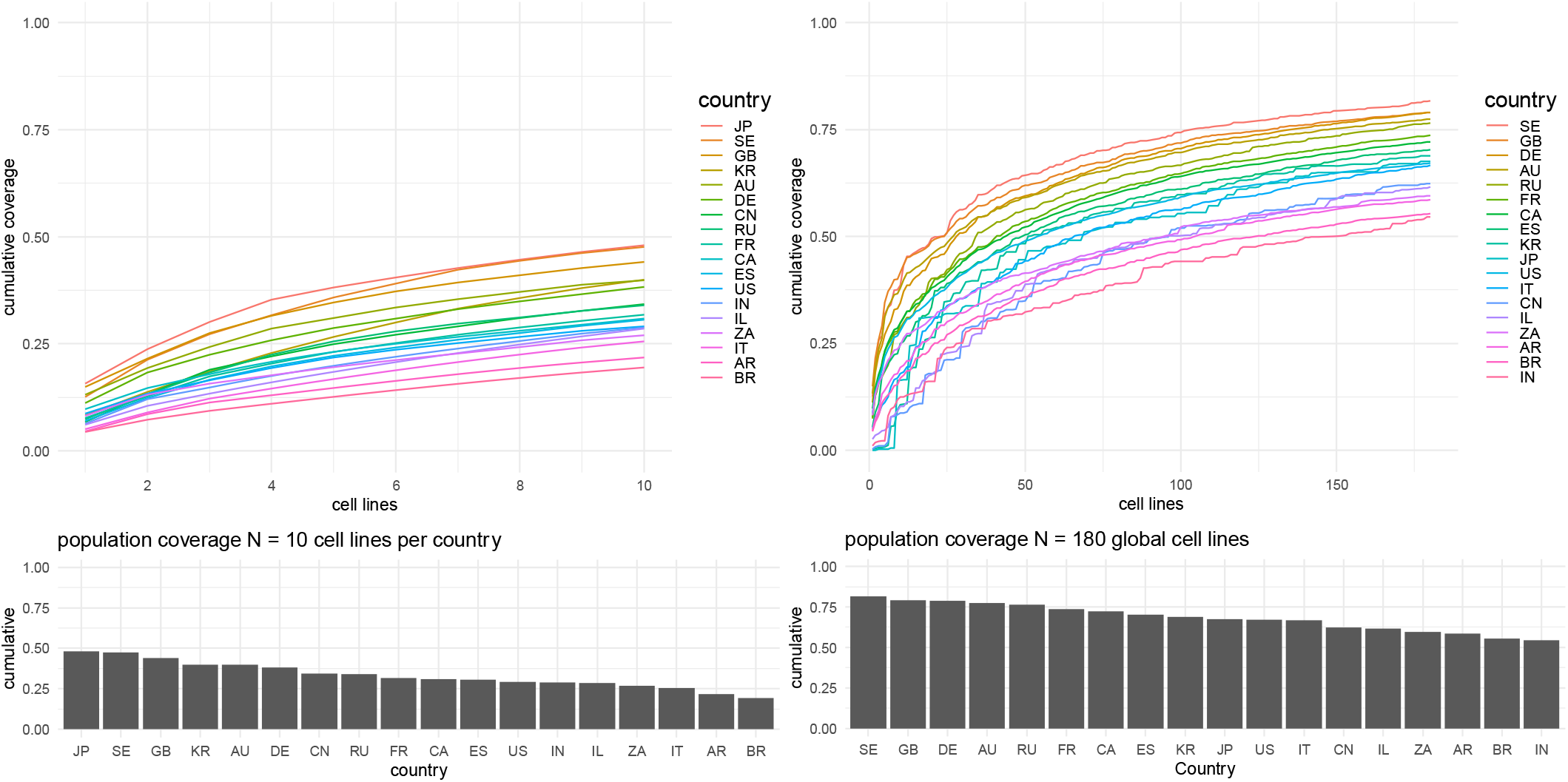
Independent coverage analysis for 18 countries for their local optimal 10 (possibly overlapping) haplotypes. Coverage ranges from 19.5% (BR) to 48.0% (JP) (left). Coverage analysis for 180 haplotypes optimized to maximize average coverage across all 18 populations in each iteration. Coverage ranges from 81.7% (SE) to 54.6% (IN) with an average coverage of 68.4%.

The advantages of global collaboration are illustrated in Figure 2 and Table 2 which compare the percentage patient coverage for local banks versus a global bank of 180 cell lines. The number 180 was chosen for the hypothetical global bank as it would require the same resource as 10 lines being developed in each of the 18 countries. A local bank of 10 local cell lines would cover 47.6% of patients in Sweden and only 19.5% of patients in Brazil. In contrast, a global bank of 180 cell lines, optimized for average coverage across all 18 countries, would increase coverage to 81.7% in Sweden and 55.3% in Brazil. On average, global collaboration and the sharing of materials more than doubles coverage (2.1x). However, the benefits of global collaboration are not uniformly distributed. Countries in North and South America, particularly those with diverse European, non-European and admixed populations (e.g., Canada, Argentina, and Brazil), experience the greatest benefit. In contrast, East and South Asian countries, such as India, Japan, China, and Korea, see a slightly lower benefit (1.7x on average) compared to a locally optimal bank of 10 cell lines. Note that the optimization is maximizing the average coverage, and each country is weighted equally.

The top 30 global coverage haplotypes are presented in Table 3, showing their coverage across each country and also with the number of donors and cord blood units that are homozygous for each haplotype, serving as the potential source of starting material across the 18 countries. The full list of the top 180 global coverage haplotypes is provided in Supplemental Material 2. Figure 3 illustrates the top 30 global haplotypes clustered by country (columns) and clustered by both country and haplotype (rows and columns). The 1D clustering by country generally aligns with continental geographic regions, with a large cluster comprising 9 countries in Europe or with predominantly European-origin population in their donor/CBU registry (SE, DE, GB, AU, CA, US, ZA, FR, RU). (Refer to Supplemental Figure 1 for a PCA analysis based on the top 30 coverage haplotypes, which resembles a world map). The next cluster includes countries from the Mediterranean region (ES, IT, IL) or countries with substantial Mediterranean-origin populations (BR, AR). The final cluster (far left) consists of four countries from Eastern and Southern Asia (JP, KR, IN, CN). The 2D clustering heatmap provides a clearer view of how specific haplotypes provide coverage for specific world regions with haplotypes ranked 9, 13 and 16 providing coverage for JP & KR and haplotypes 7, 18, 22 and 24 providing coverage for KR, IN and CN. Haplotypes 6 and 22 provide specific benefit to India with haplotype 6 (A*01:01∼B*57:01∼DRB1*07:01) providing some coverage to European-origin populations and haplotype 22 (A*33:03∼B*44:03∼DRB1*07:01) providing coverage for Korean patients.

**Table 3.**
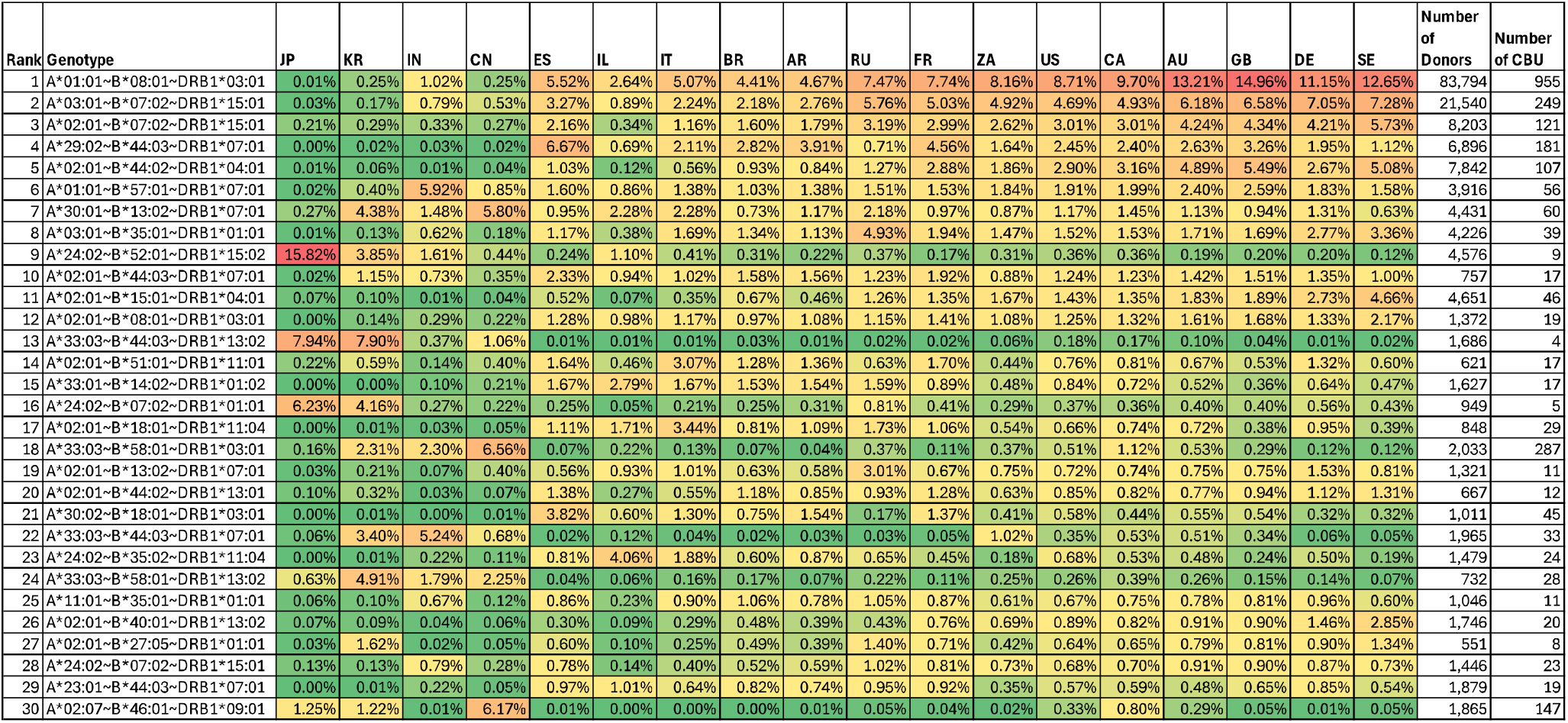
Top 30 global optimal homozygous haplotypes, the marginal coverage (percentage) this haplotype represents in each country, and the number of donors and cord blood units (CBU) in the 18 countries identified as being homozygous for these haplotypes. The columns for the 18 countries are ordered based on clustering by similarity. The color scale ranges from red (high) to low (green).

**Figure 3.**
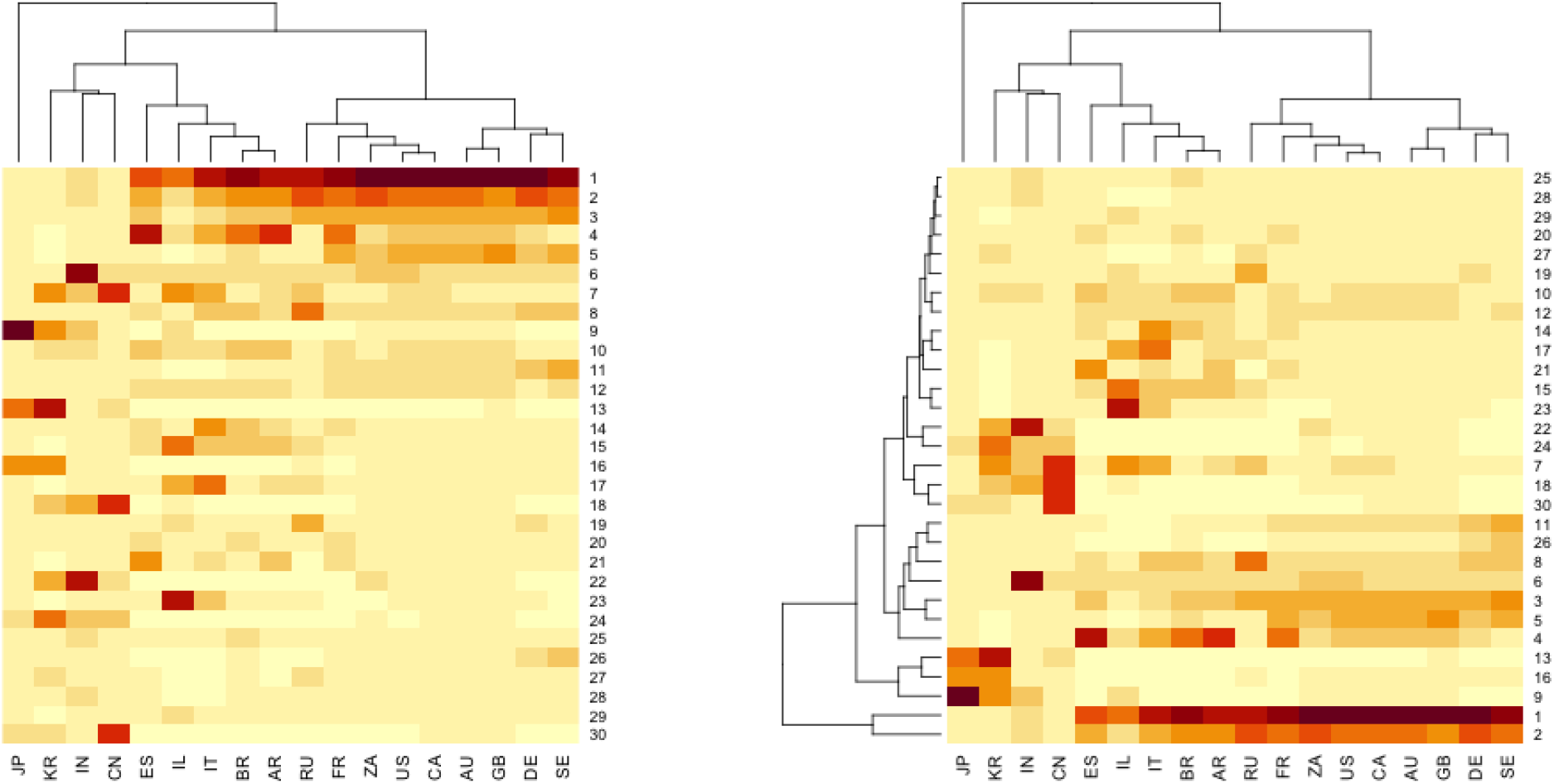
1D and 2D clustering of the top 30 global haplotypes.

## Discussion

Treatments derived from pluripotent stem cells have started to move into clinical use (Sugita et al. 2020). The issue of mitigating immune reactivity against allogeneic cells has been addressed by either looking to genetically alter stem cell lines to reduce the potential for immune cell activation (Han et al. 2019), or by using HLA matched donor cells (Sullivan, Fairchild, et al. 2020). This latter approach requires the identification of individuals who have specific HLA types, with most studies focusing on donors who have two copies of high frequency HLA haplotypes in the target population, so that immune matching in the host versus graft direction (HvG) is achieved for patients carrying one copy of that haplotype (Gourraud et al. 2012). Given the global allelic variation in HLA genes, countries looking to develop ‘haplobanks’ of donor iPSC lines have undertaken analysis of local HLA genotype data to identify the most frequent HLA haplotypes in their own populations (Roh et al. 2020; Nakatsuji, Nakajima, and Tokunaga 2008; Taylor et al. 2012; Lee et al. 2018; Belen Alvarez-Palomo et al. 2022; Martins De Oliveira et al. 2023). Our results are comparable to these studies, although many of them only consider low-resolution HLA due to ambiguity in the registry typing. In some countries this work has gone beyond a theoretical analysis and haplobanks of iPSC lines have been created for clinical and research use (Yoshida et al. 2023; Martins De Oliveira et al. 2023).

In this analysis, as expected and seen in many previous studies cited above, there was a range of patient matching coverage when the top 10 or 30 haplotypes were identified in each donor pool and used to match against that population. Countries with lower coverage tend to be those with considerable admixture (South Africa, US, Canada) whereas the countries with higher coverage are island nations (Japan, UK) or countries with otherwise low admixture (Sweden, Korea). This analysis illustrates the way that collaboration may benefit nascent haplobanks and avoid redundancy of effort in creating the same lines in different countries.

An argument can be made that the resources required for all 18 countries used in this analysis to identify and bank cell lines from homozygous donors that match their top 10 HLA haplotypes is equivalent to a global effort to identify 180 donors optimized to cover all populations. Using all 18 countries’ data in a global analysis enabled the identification of a top 180 of haplotypes that give a range of coverage of 81.7% in Sweden to 54.6% in India with an average of 68.4%. For all countries, an optimized global 180 haplotypes yields greater coverage than the local 30 haplotypes, but with significantly less overall investment and resource (Table 2). This illustrates the benefits that can be realized by working collaboratively across countries which are developing iPSC derived therapies to increase coverage across all populations.

A further consideration prior to the development of clinical iPSC banks is the source of donor material. Analyses such as this indicate the optimum homozygous HLA genotypes to target but raise questions about how to procure starting material. Making use of unrelated donor hematopoietic stem cell registries and cord blood banks prevents the need for more HLA typing of populations. As shown in Table 3, for the top 30 global haplotypes the relative likelihood of identifying suitable donors for a given haplotype from stem cell registries or cord banks can be established, considering that the number of ultimately suitable donors will be further decreased by giving preference to blood group O and female donors to limit the impact of blood group and minor histocompatibility antigen reactivity respectively. Both sources of donor material present issues such as informed consent from the donor, compensation for the use of their cells and ownership of future intellectual property from therapies derived from an individual’s cells (de Rham & Villard 2014), although these have been overcome in some countries allowing the development of banks for clinical and research use. Homozygosity at A, B and DRB1 is rare and may not be necessary to achieve the desired histocompatibility (Rushakoff et al. 2022). One way to reduce the limitation in starting material would be to allow genotypes that are partially homozygous with the hope of finding a balance between reducing the degree of mismatch in the rejection direction while increasing the availability of starting material. In future models, special attention should also be paid to patient NK receptors to try to avoid alloreactivity against donor derived cells which might lack the appropriate HLA ligands to provide ‘self’ inhibitory signals (Ichise et al. 2017).

### From modeling to bank building

In Japan a clinical grade HLA haplobank has been developed by recruiting donors from pools of platelet, hematopoietic stem cell and cord donors (Yoshida S et al, 2023). After receiving an invitation to participate, the onus was on the donors to contact the coordinators to initiate the consent process. In Australia, the potential for creating a haplobank using cord blood from the government funded BMDI Cord Blood Bank is being explored, with the local top 18 donors with suitable HLA haplotypes selected (Abberton KM et al. 2022). In Europe, ongoing haplobanking efforts emphasize the use of cord donors as the most accessible source of cells, especially where cord units have a nucleated cell count too low to be viable for clinical transplantation (Belen Alvarez-Palomo et al. 2022). As haplobanking in different countries continues to develop it is hoped that global collaboration can confer the maximum benefit to patients in relation to HLA matching.

The optimal model for iPSC banking, whether public or private, remains a topic of debate. While adult donor HSC registries follow a public model, cord blood banking has utilized public, private, and hybrid approaches. Notably, iPSC banks (Abberton et al. 2022) are currently managed by both public and private entities. Aligning iPSC banking with public models would resonate with the principles of organizations such as WMDA and GAiT.

Although the model proposed on this paper is optimized to provide the highest *average* patient coverage across countries, it would eventually need to be extended to incorporate equity between countries. Specifically, countries with predominantly European ancestry tend to benefit the most from sharing under the current models and have higher patient coverage than Asian countries. The model doesn’t consider heterogeneity within a country. For a country like the US with a European majority, the approach used here will not provide equity within all subpopulations. This can be addressed by modeling sub-populations within countries explicitly, which in turn requires additional data collection and standardization. Yet another equity challenge is the fact that the 18 countries included in the model have different population sizes and different levels of access to treatment. The goal of providing coverage for the highest percentage of patients must be balanced against the goal of providing equity to patients.

Furthermore, the use of donor and cord blood unit HLAs as a model for the HLA of the patient population is a limitation to the extent that there are HLA types associated with diseases treatable by iPSC-based therapies or more generally if there is bias in how donors and cord blood units are recruited and banked that may impose a different HLA distribution compared to patients.

There are many practical limitations to global sharing. For example, cord blood has declined in use as a source of HSC transplants, which seemingly would make the use of these units as starting material for an iPSC haplobank particularly attractive, in particular because of how readily these units are available.

However, the 9000 cord blood units in Japan (Watanabe-Okochi et al. 2024) are not available for export. Furthermore, many inventoried cord blood units may not be consented for use as starting material for an iPSC bank, and re-consenting may be difficult or impossible. In addition, the use of cord blood exclusively will have challenges due to the lower numbers of donors. Another option is to source cells from adult donors, but consenting or reconsenting adult donors can be challenging. A new public iPSC bank in Brazil (Martins De Oliveira et al. 2023) contacted members of the Brazilian bone marrow registry (REDOME) and found that less than half of the registry members agreed to provide cells for commercial use (personal communication, Danielli Oliveira). One might expect a higher response rate for donation to a public iPSC bank compared with a private one. Although some organizations have pursued financial renumeration to motivate donation of cellular material, the WMDA has outlined several ethical concerns related to renumeration of donors for cell and gene therapy development (Hamad et al. 2024) and advises non-renumeration for the safety and well-being of both the donor and the patient.

In summary, the data presented illustrates the patient HLA matching coverage benefits of iPSC banks operating at a global scale, thereby reducing the redundancy of iPSC line production. Such collaboration would enable the treatment of more patients with iPSC-derived therapies while minimizing overall investment. While this analysis is based on several assumptions, including the feasibility of sharing and transferring iPSC lines across regulatory jurisdictions, where production comparability would need to be established, it nonetheless serves as a proof of principle. We believe this approach signposts a path forward for optimizing iPSC therapies across the broad expanse of HLA polymorphism.

## Abbreviations

iPSC: Induced Pluripotent Stem Cell
HLA: Human Leukocyte Antigen
CBU: Cord Blood Unit
GAiT: Global Alliance for iPSC Therapy
ESC: Embryonic Stem Cell
HSC: Hematopoietic Stem Cell
NK: Natural Killer (Cell)
WMDA: World Marrow Donor Association
REDOME: Registro Brasileiro de Doadores Voluntários de Medula Óssea (Brazilian Registry of Voluntary Bone Marrow Donors)
BMDI: Bone Marrow Donor Institute
AR: Argentina
AU: Australia
BR: Brazil
CA: Canada
CN: China
DE: Germany
ES: Spain
FR: France
GB: UK
IL: Israel
IN: India
IT: Italy
JP: Japan
KR: S. Korea
RU: Russia
SE: Sweden
US: USA
ZA: South Africa

## Acknowledgements

This work was supported by the Global Alliance for iPSC Therapy (GAiT). MM is supported by a grant from the US Office of Naval Research (N00014-23-1-2057). The authors wish to acknowledge and thank Dennis Confer, Mike Boo, Kaveh Mirfakhraie, Emile Nuwaysir, Michelle Hunt, Amanda Mack, Derek Hei, Jamie Thompson, Marcelo Fernandez-Viña, Malek Kamoun, Craig Taylor, Danielli Oliveira, Luís Cristóvão Pôrto, Lygia V Pereira, Antonio Carlos Campos de Carvalho, Loren Gragert, Yoram Louzoun and Pierre-Antoine Gourraud for thoughtful discussions about the study. We wish to thank Alicia Venter from the World Marrow Donor Program for facilitating access to the data. Michael Wright performed proofreading and provided editorial input to the manuscript.

## Supplemental Material

**Supplemental Material 1**. Spreadsheet with optimal single ground coverage models for top 30 haplotypes. Note that for RU and ZA the list is less than 30 due to the low sample size.

**Supplemental Material 2**. The optimal 180 global coverage homozygous genotypes with counts of donors and CBU by country of origin with the corresponding genotype.

**Supplemental Figure 1**. Principal Components Analysis (PCA) of top 100 haplotypes

## References

Álvarez-Palomo, Belén, Iris García-Martinez, Jorge Gayoso, Angel Raya, Anna Veiga, María Luisa Abad, Adolfo Eiras, et al. 2021. “Evaluation of the Spanish Population Coverage of a Prospective HLA Haplobank of Induced Pluripotent Stem Cells.” Stem Cell Research and Therapy 12 (1). 10.1186/s13287-021-02301-0.

Belen Alvarez-Palomo, Anna Veiga, Angel Raya, Margarita Codinach, Silvia Torrents, Laura Ponce Verdugo, Clara Rodriguez-Aierbe, et al. 2022. “Public Cord Blood Banks as a Source of Starting Material for Clinical Grade HLA-Homozygous Induced Pluripotent Stem Cells.” Stem Cell Research & Therapy 13 (1). 10.1186/s13287-022-02961-6.

Crow, Diana. 2019. “Could iPSCs Enable ‘Off-the-Shelf’ Cell Therapy?” Cell 177 (7): 1667–69. 10.1016/j.cell.2019.05.043.

Doss, Michael Xavier, and Agapios Sachinidis. 2019. “Current Challenges of iPSC-Based Disease Modeling and Therapeutic Implications.” Cells 8 (5): 403. 10.3390/cells8050403.

Fairchild, Paul J, Christopher Horton, Priyoshi Lahiri, Kumaran Shanmugarajah, and Timothy J Davies. 2016. “Beneath the Sword of Damocles: Regenerative Medicine and the Shadow of Immunogenicity.” Regenerative Medicine 11 (8): 817–29. 10.2217/rme-2016-0134.

Gourraud, Pierre-Antoine, Gilson Leena, Mathilde Girard, and Marc Peschanski. 2012. “The Role of Human Leukocyte Antigen Matching in the Development of Multiethnic ‘Haplobank’ of Induced Pluripotent Stem Cell Lines.” Stem Cells (Dayton, Ohio) 30 (2): 180–86. 10.1002/stem.772.

Hamad, Lina, Salmah Mahmood Ahmed, Eefke van Eerden, Suzanna M. van Walraven, World Marrow Donor Association Cellular Therapy Committee, and Laura Machin. 2024. “Remuneration of Donors for Cell and Gene Therapies: An Update on the Principles and Perspective of the World Marrow Donor Association.” Bone Marrow Transplantation 59 (5): 580–86. 10.1038/s41409-024-02246-x.

Han, Xiao, Mengning Wang, Songwei Duan, Paul J. Franco, Jennifer Hyoje-Ryu Kenty, Preston Hedrick, Yulei Xia, et al. 2019. “Generation of Hypoimmunogenic Human Pluripotent Stem Cells.” Proceedings of the National Academy of Sciences of the United States of America 116 (21): 10441–46. 10.1073/pnas.1902566116.

Ichise, Hiroshi, Seiji Nagano, Takuya Maeda, Masaki Miyazaki, Yuki Miyazaki, Hiroto Kojima, Nobuyo Yawata, et al. 2017. “NK Cell Alloreactivity against KIR-Ligand-Mismatched HLA-Haploidentical Tissue Derived from HLA Haplotype-Homozygous iPSCs.” Stem Cell Reports 9 (3): 853–67. 10.1016/j.stemcr.2017.07.020.

Israeli, Sapir, Loren Gragert, Abeer Madbouly, Pradeep Bashyal, Joel Schneider, Martin Maiers, and Yoram Louzoun. 2023. “Combined Imputation of HLA Genotype and Self-Identified Race Leads to Better Donor-Recipient Matching.” Human Immunology 84 (12): 110721. 10.1016/j.humimm.2023.110721.

Israeli, Sapir, Loren Gragert, Martin Maiers, and Yoram Louzoun. 2021. “HLA Haplotype Frequency Estimation for Heterogeneous Populations Using a Graph-Based Imputation Algorithm.” Human Immunology 82 (10): 746–57. 10.1016/j.humimm.2021.07.001.

Israeli, Sapir, Elizabeth F. Krakow, Martin Maiers, Corinne Summers, and Yoram Louzoun. 2023. “Trans-Population Graph-Based Coverage Optimization of Allogeneic Cellular Therapy.” Frontiers in Immunology 14 (May):1069749. 10.3389/fimmu.2023.1069749.

Jonna Clancy, Kati Hyvärinen, Jarmo Ritari, Tiina Wahlfors, Jukka Partanen, and Satu Koskela. 2022. “Blood Donor Biobank and HLA Imputation as a Resource for HLA Homozygous Cells for Therapeutic and Research Use.” Stem Cell Research & Therapy 13 (1). 10.1186/s13287-022-03182-7.

Kawamura, Takuji, Shigeru Miyagawa, Satsuki Fukushima, Akira Maeda, Noriyuki Kashiyama, Ai Kawamura, Kenji Miki, et al. 2016. “Cardiomyocytes Derived from MHC-Homozygous Induced Pluripotent Stem Cells Exhibit Reduced Allogeneic Immunogenicity in MHC-Matched Non-Human Primates.” Stem Cell Reports 6 (3): 312–20. 10.1016/j.stemcr.2016.01.012.

Kobold, Sabine, Nils Bultjer, Glyn Stacey, Sabine C. Mueller, Andreas Kurtz, and Nancy Mah. 2023. “History and Current Status of Clinical Studies Using Human Pluripotent Stem Cells.” Stem Cell Reports 18 (8): 1592–98. 10.1016/j.stemcr.2023.03.005.

Lee, Suji, Ji Young Huh, David M. Turner, Soohyeon Lee, James Robinson, Jeremy E. Stein, Sung Han Shim, et al. 2018. “Repurposing the Cord Blood Bank for Haplobanking of HLA-Homozygous iPSCs and Their Usefulness to Multiple Populations.” Stem Cells (Dayton, Ohio) 36 (10): 1552–66. 10.1002/stem.2865.

Martins De Oliveira, Marcio Lassance, Bernardo Rangel Tura, Mauro Meira Leite, Eduardo José Melo Dos Santos, Luís Cristóvão Pôrto, Lygia V. Pereira, and Antonio Carlos Campos De Carvalho. 2023. “Creating an HLA-Homozygous iPS Cell Bank for the Brazilian Population: Challenges and Opportunities.” Stem Cell Reports 18 (10): 1905–12. 10.1016/j.stemcr.2023.09.001.

Morizane, Asuka, Tetsuhiro Kikuchi, Takuya Hayashi, Hiroshi Mizuma, Sayuki Takara, Hisashi Doi, Aya Mawatari, et al. 2017. “MHC Matching Improves Engraftment of iPSC-Derived Neurons in Non-Human Primates.” Nature Communications 8 (1): 1–12. 10.1038/s41467-017-00926-5.

Nakatsuji, Norio, Fumiaki Nakajima, and Katsushi Tokunaga. 2008. “HLA-Haplotype Banking and iPS Cells.” Nature Biotechnology 26 (7): 739–40. 10.1038/nbt0708-739.

Pappas, Derek James, Pierre-Antoine Gourraud, Caroline Le Gall, Julie Laurent, J. Laurent, Alan O Trounson, Natalie DeWitt, and Sohel Talib. 2015. “Proceedings: Human Leukocyte Antigen Haplo-Homozygous Induced Pluripotent Stem Cell Haplobank Modeled After the California Population: Evaluating Matching in a Multiethnic and Admixed Population.” Stem Cells Translational Medicine 4 (5): 413–18. 10.5966/sctm.2015-0052.

Petrus-Reurer, Sandra, Marco Romano, Sarah Howlett, Joanne Louise Jones, Giovanna Lombardi, and Kourosh Saeb-Parsy. 2021. “Immunological Considerations and Challenges for Regenerative Cellular Therapies.” Communications Biology 4 (1): 798. 10.1038/s42003-021-02237-4.

Roh, Eun Youn, Sohee Oh, Sohee Oh, Jong Hyun Yoon, Byoung Jae Kim, Byoung Jae Kim, Eun Young Song, and Sue Shin. 2020. “Umbilical Cord Blood Units Cryopreserved in the Public Cord Blood Bank: A Breakthrough in iPSC Haplobanking?” Cell Transplantation 29 (July):963689720926151. 10.1177/0963689720926151.

Rushakoff, Joshua A., Loren Gragert, Marcelo J. Pando, Darren Stewart, Edmund Huang, Irene Kim, Stanley Jordan, et al. 2022. “HLA Homozygosity and Likelihood of Sensitization in Kidney Transplant Candidates.” Transplantation Direct 8 (5): e1312. 10.1097/TXD.0000000000001312.

Solomon, Susan, Fernando Pitossi, and Mahendra S Rao. 2015. “Banking on iPSC–Is It Doable and Is It Worthwhile.” Stem Cell Reviews 11 (1): 1–10. 10.1007/s12015-014-9574-4.

Sugita, Sunao, Michiko Mandai, Yasuhiko Hirami, Seiji Takagi, and Tadao Maeda. 2020. “HLA-Matched Allogeneic iPS Cells-Derived RPE Transplantation for Macular Degeneration.”

Sullivan, Stephen, Paul J. Fairchild, Steven G.E. Marsh, Carlheinz Müller, Marc L. Turner, Marc Turner, Jihwan Song, and David Turner. 2020. “Haplobanking Induced Pluripotent Stem Cells for Clinical Use.” Stem Cell Research 49:102035. 10.1016/j.scr.2020.102035.

Sullivan, Stephen, Patrick Ginty, Siofradh McMahon, Michael May, Susan L. Solomon, Andreas Kurtz, Glyn N. Stacey, et al. 2020. “The Global Alliance for iPSC Therapies (GAiT).” Stem Cell Research 49 (December):102036. 10.1016/j.scr.2020.102036.

Taylor, Craig J, Eleanor M Bolton, Susan Pocock, Linda D Sharples, Roger A Pedersen, and J Andrew Bradley. 2005. “Banking on Human Embryonic Stem Cells: Estimating the Number of Donor Cell Lines Needed for HLA Matching.” Lancet 366 (9502): 2019–25. 10.1016/S0140-6736(05)67813-0.

Taylor, Craig J, Sarah Peacock, Afzal N Chaudhry, J Andrew Bradley, and Eleanor M Bolton. 2012. “Generating an iPSC Bank for HLA-Matched Tissue Transplantation Based on Known Donor and Recipient HLA Types.” Cell Stem Cell 11 (2): 147–52. 10.1016/j.stem.2012.07.014.

Tian, Pei, Andrew Elefanty, Edouard G. Stanley, Jennifer C. Durnall, Lachlan H. Thompson, and Ngaire J. Elwood. 2022. “Creation of GMP-Compliant iPSCs From Banked Umbilical Cord Blood.” Frontiers in Cell and Developmental Biology 10:835321. 10.3389/fcell.2022.835321.

Turner, Marc, Stephen Leslie, Nicholas G. Martin, Marc Peschanski, Mahendra Rao, Craig J. Taylor, Alan Trounson, David Turner, Shinya Yamanaka, and Ian Wilmut. 2013. “Toward the Development of a Global Induced Pluripotent Stem Cell Library.” Cell Stem Cell 13 (4): 382–84. 10.1016/j.stem.2013.08.003.

Watanabe-Okochi, Naoko, Takeshi Odajima, Miyuki Ito, Naoya Yamada, Manami Shinozaki, Mutsuko Minemoto, Fumihiko Ishimaru, Kazuo Muroi, and Minoko Takanashi. 2024. “Criteria for Storage of Cord Blood Units at Japan’s Largest Cord Blood Bank.” Vox Sanguinis 119 (8): 867–77. 10.1111/vox.13687.

Wilmut, Ian, Stephen Leslie, Nicholas G. Martin, Marc Peschanski, Mahendra S. Rao, Alan O Trounson, David M. Turner, et al. 2015. “Development of a Global Network of Induced Pluripotent Stem Cell Haplobanks.” Regenerative Medicine 10 (3): 235–38. 10.2217/rme.15.1.

Xu, Huaigeng, Bo Wang, Miyuki Ono, Akihiro Kagita, Kaho Fujii, Noriko Sasakawa, Tatsuki Ueda, et al. 2019. “Targeted Disruption of HLA Genes via CRISPR-Cas9 Generates iPSCs with Enhanced Immune Compatibility.” Cell Stem Cell 24 (4). 10.1016/j.stem.2019.02.005.

Yoshida, Shinsuke, Tomoaki M. Kato, Yoshiko Sato, Masafumi Umekage, Tomoko Ichisaka, Masayoshi Tsukahara, Naoko Takasu, and Shinya Yamanaka. 2023. “A Clinical-Grade HLA Haplobank of Human Induced Pluripotent Stem Cells Matching Approximately 40% of the Japanese Population.” Med (New York, N.Y.) 4 (1): 51-66.e10. 10.1016/j.medj.2022.10.003.

