## Supplementary material for "Harnessing Global HLA Data for Enhanced Patient Matching in iPSC Haplobanks": Supplemental Figure 1.docx

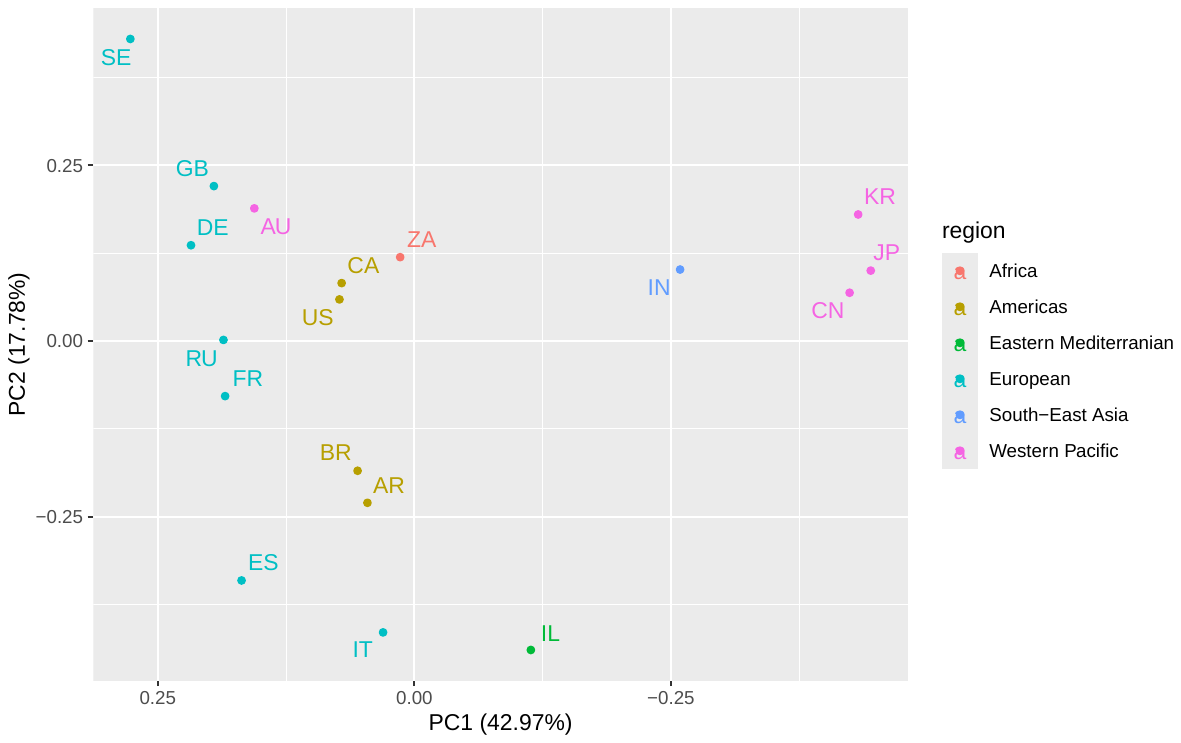
Supplemental Figure 1 - Principal components analysis of the coverage proportion for the top 30 global coverage genotypes. The colors reflect the corresponding world region according to the World Health Organization (who.org)
